# Direct measurement and reconstruction of intact polyclonal IgG repertoires to preserve molecular and functional connectivity

**DOI:** 10.64898/2026.07.30.741668

**Authors:** Thomas Holmark, Despoina Mavridou, Ziran Zhai, Annika A.M. van der Zon, Constantin Blöchl, Elena Domínguez-Vega, Andrea F.G. Gargano

**Affiliations:** Analytical Chemistry Group, Van ’t Hoff Institute for Molecular Sciences, University of Amsterdam, Science Park 904, 1098XH, Amsterdam, the Netherlands; Centre for Analytical Sciences Amsterdam (CASA), Amsterdam, the Netherlands; Center for Proteomics and Metabolomics, Leiden University Medical Center, Albinusdreef 2, 2333ZA, Leiden, the Netherlands

## Abstract

Antibody function depends on the molecular pairing of antigen-binding Fab regions with Fc domains, linking antigen specificity to effector potential. However, antibody repertoire methods analyze these subunits separately, creating an inference problem in which Fc-Fab connectivity is lost. Here, we introduce a multi-level HPLC-MS workflow integrating native nanoflow cation-exchange chromatography-MS (nCEC-MS) of intact serum IgG with middle-up Fab and Fc/2 profiling. Native nCEC-MS reduces spectral congestion compared with denaturing reversed-phase HPLC-MS, enabling detection of 56-74 intact IgG mass features from 2 μg of purified IgG per donor. Middle-up analysis provided complementary information on Fab diversity, subclass, and allotype, revealing donor-specific profiles. Reconstructing intact IgG1 masses from independently measured Fab and Fc/2 subunits partially recovers Fc-Fab molecular connectivity and supports interpretation of native intact mass features. Together, this framework provides an integrated view of the human IgG repertoire and establishes a foundation for studying its molecular topology in health and disease.

## Introduction

The immune system produces immunoglobulins (Igs) to recognize pathogens, malignant cells, and other immune challenges. The effectiveness of immune responses depends strongly on each individual’s unique repertoire of circulating antibodies and immune cells. Among human plasma immunoglobulin isotypes, IgG is the most abundant and comprises four subclasses, IgG1–4, which are defined by distinct amino acid sequences, structural properties, and effector functions.^1,2^

Circulating IgGs exhibit extraordinary structural variability driven by clonal selection and somatic hypermutation. IgG diversity is concentrated in the variable domains of both the heavy and light chains, each of which contains three complementarity-determining regions (CDRs): CDR1, CDR2, and CDR3. This structural complexity underlies a diverse repertoire of IgG clones circulating in human plasma.^3,4^ IgGs mediate immunological responses by binding antigens through their Fab regions, while their Fc regions engage downstream immune receptors, including Fcγ receptors, to activate distinct effector pathways. Beyond direct antigen-mediated activation, B-cell differentiation is further shaped by secondary signals, including those from pattern recognition receptors.^5^ Thus, IgG function is determined mainly by Fc-mediated engagement of immune effector systems, including Fcγ receptors and the complement cascade, whereas the Fab domain is critical in antigen-mediated recognition.^6^

To characterize the IgG repertoire, the field has advanced beyond conventional bottom-up proteomics, which resolves antibodies primarily at the peptide level, towards middle-up and middle-down mass spectrometry strategies. Rather than reducing antibodies to short peptides, these approaches retain larger antibody structural units by combining hinge-region enzymatic cleavage, such as IdeS or IgdE digestion, with increasingly capable HPLC-MS and charge-detection mass spectrometry platforms.^5,7–12^ Current approaches can be broadly categorized into three strategies: (i) Fab profiling, as introduced by Bondt et al., which isolates the IgG1 Fab domain to track clonal responses and sequence variability;^10,11,13,14^ (ii) Fc profiling, as developed by Blöchl et al., which focuses on the Fc/2 subunit to map global N-glycosylation, subclass, and allotype distributions;^8,15,16^ and (iii) single-ion charge detection mass spectrometry (CDMS), in which Melani et al. use CDMS to measure mass distributions of isolated heavy and light chains.^5,12,17^

These state-of-the-art middle-up strategies have yielded valuable insights into diversification of antibody responses and maturation in scenarios such as rheumatoid arthritis, COVID-19, and sepsis.^10,11,13,15,17,18^ However, they share an important limitation: they disrupt the molecular connectivity between the antigen-binding Fab region and the functional Fc region of the same intact IgG molecule. Here, we refer to this physical linkage as IgG molecular topology. This information is crucial for interpreting IgG-mediated immunity, particularly in complex IgG responses in which antigen recognition, clonal expansion, subclass usage, and Fc effector functions may be tightly coupled. Current middle-up workflows rely on offline enzymatic hydrolysis prior to analysis. Hinge-region digestion is often performed directly on the beads used for antibody enrichment, thereby separating and isolating the Fab and Fc domains before measurement. This creates a correlation gap, in which the original Fc-Fab pairings must be inferred rather than directly observed. As a result, even when antigen-specific fractions show distinct mass profiles, current middle-up analytics cannot determine whether these profiles are carried by antibodies with a particular pro-inflammatory Fc subclass on the same molecular backbone.

Approaches that combine the molecular specificity of middle-up profiling with intact-level measurements to reconstruct these domain pairings remain unavailable. Resolving IgG molecular topology requires analytical methods capable of measuring intact polyclonal IgG distributions while retaining sufficient sensitivity, mass resolution, and separation selectivity to distinguish closely related antibody proteoforms. This remains a major challenge because the circulating IgG repertoire is highly heterogeneous, spanning a broad range of clonal abundances and molecular masses. As a consequence, direct infusion approaches are not suitable for this purpose, as the high heterogeneity of the repertoire causes severe spectral congestion during ionization. Hundreds of overlapping charge-state envelopes collapse into convoluted spectra in which individual clonal masses cannot be distinguished. To measure IgG masses across a wide concentration range, HPLC separations are therefore needed to resolve intact IgG proteoforms prior to MS analysis.

Reversed-phase liquid chromatography (RP-HPLC) coupled to MS is widely used for robust protein analysis in top-down proteomics. However, the acidic organic mobile phases typically employed in RPLC unfold the antibody structure, distributing the IgG MS signal over many highly charged states. For highly heterogeneous polyclonal IgG samples, these broad charge-state distributions can increase spectral congestion and complicate the extraction and assignment of individual intact mass features, particularly when multiple species co-elute. Native nanoflow HPLC-MS provides a complementary approach by maintaining proteins under non-denaturing conditions during separation and ionization. Unlike denaturing RPLC-MS, native HPLC-MS methods using buffered systems near physiological pH preserve non-denatured protein structures during analysis. The resulting compact charge-state distributions and wider *m/z* spacing between adjacent charge states reduce spectral overlap and improve the interpretability of complex intact IgG spectra. These characteristics make native MS particularly advantageous for preserving and examining intact-level molecular information within polyclonal IgG repertoires.

In this work, we introduce a multi-level analytical framework that integrates high-resolution native nanoflow cation-exchange chromatography (nCEC) coupled to MS^19^ of intact polyclonal IgG with parallel middle-up Fc/2 and Fab subunit profiling.^10^ By aligning intact native IgG distributions with isolated Fab and Fc/2 subunit profiles, we present what we believe is the first direct method for reconstructing clonal human IgG molecular topologies obtained from individuals. This workflow provides a route to connect clonal mass profiles, Fab diversity, subclass, allotype, and Fc glycosylation information within the same analytical framework. Preserving and integrating this molecular context enables a more complete view of the human IgG repertoire and opens new frontiers in structural immunology.

## Results

### Native nCEC-MS resolves intact human IgG proteoform repertoires

Current state-of-the-art HPLC-MS(/MS) methods for clone profiling and detailed characterization of endogenous immunoglobulins frequently rely on reversed-phase liquid chromatography (RPLC) coupled to MS. While these methods provide high sensitivity and chemical selectivity for isolated antibody fragments or engineered monoclonal therapies,^5,12,20,21^ they offer limited separation power when applied to fully intact polyclonal IgG mixtures. Moreover, the acidic and organic conditions typically used in RPLC unfold antibody structures, distributing the IgG signal across multiple highly charged states and increasing spectral complexity.

To evaluate alternative strategies for intact IgG analysis, we compared denaturing nRPLC-MS with native nCEC-MS using monoclonal antibodies (mAbs) and serum-derived IgGs from donor M54 (Supplementary Figures 1 and 2). As shown in Supplementary Figure 1A, RPLC provided limited selectivity for the selected mAbs, resulting in co-elution of the three mAbs. By contrast, native nCEC-MS achieved charge-based separation under near-native mobile phase conditions using volatile ammonium acetate buffers with a pH range from 5.0 (mobile phase A) to 8.5 (mobile phase B, Supplementary Figure 1B). This resulted in a broader separation window for the three mAb species and their charge variants. The resolution between antibody species with conventional RPLC-MS has been demonstrated, however, the concentrations required are not feasible for serum derived IgGs.

When applied to endogenous serum IgG from donor M54, RPLC-MS generated broad and overlapping elution profiles, where identical, unresolved mass spectra in the *m/z* 2000–3000 were observed continuously over an elution window of 3 to 4 minutes (Supplementary Fig. 2A). Native nCEC-MS, however, preserved compact IgG charge-state envelopes in the *m/z* 5000–7000 range, reducing spectral overlap and enabling high-quality mass spectra to be extracted from 0.2 min chromatographic slices (Supplementary Figure 2B).

Having established the native nCEC-MS workflow, it was applied to polyclonal IgG isolated from three healthy serum donors and combined intact-level measurements with middle-up Fc/2 and Fab subunit profiling, as visualized in Figure 1. IgGs were purified from individual sera using affinity chromatography beads that bind to the Fc of all IgG subclasses. As a result, intact IgG profiles capture the full spectrum of serum IgGs sequence variants in the sample (i.e. clones, subclasses and allotypes). For intact analysis, samples were treated with EndoS2 to hydrolyze specifically Fc N-glycans while preserving the underlying multi-domain antibody assembly. Deglycosylation efficiency exceeded 95% across all donors, as detailed in Supplementary Figure 3. The final concentration of each sample was estimated to be approximately 1 mg/mL.

**Fig. 1:**
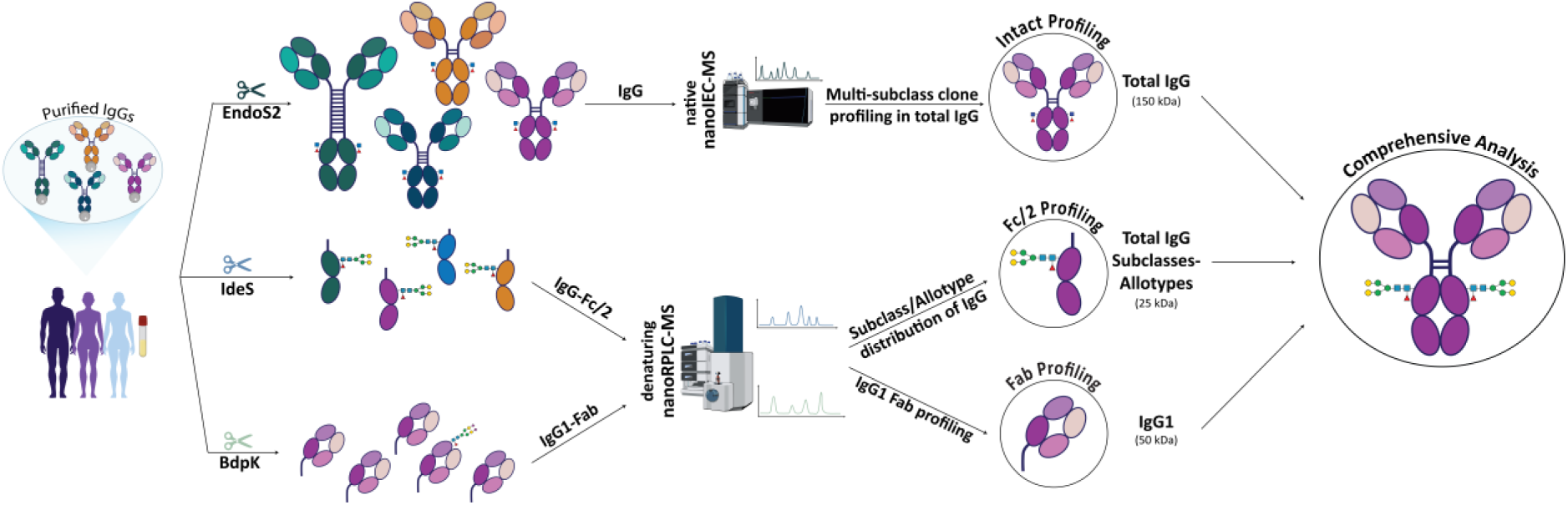
Multi-level HPLC-MS workflow for intact and subunit-level profiling of serum IgG. Serum IgGs were purified from individual donors and processed in parallel for native intact analysis and middle-up subunit profiling. For intact analysis, EndoS2 digestion was used to reduce Fc N-glycan heterogeneity while preserving the multi-domain antibody assembly, followed by native nCEC-MS. For middle-up analysis, IdeS and BdpK digestion generated Fc/2 and IgG1 Fab subunits, respectively, which were analyzed by nRPLC-MS. Integration of intact IgG mass features with Fab and Fc/2 information enables multi-level characterization of donor-specific IgG repertoires.

Figure 2A shows the 20 min native nCEC-MS separation of donor M54. Across the three donors, we detected 74 abundant intact IgG mass features for donor M54, 56 for donor F42, and 67 for donor F55. For donor M54, the ten most abundant mass features, based on deconvoluted mass intensities, were selected and visualized as extracted ion chromatograms (EICs). The corresponding deconvoluted mass spectra are shown in matching colors according to their retention times. The TICs, EICs, and corresponding deconvoluted mass spectra for donors F42 and F55 are presented in Supplementary Figure 4.

**Fig. 2:**
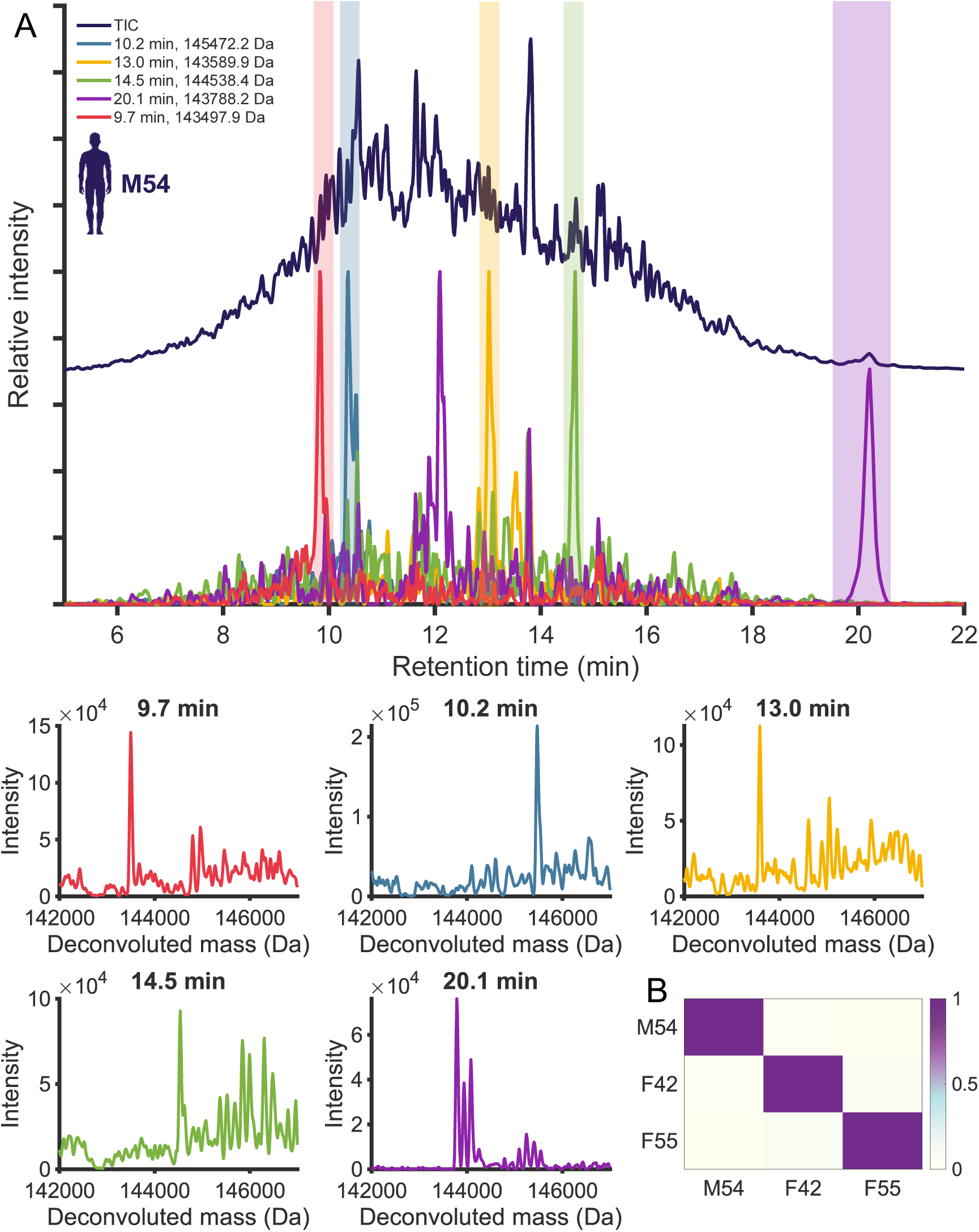
Native nCEC-MS resolves donor-specific intact IgG proteoform repertoires. (A) Total ion chromatogram of donor M54 acquired by native nCEC-MS, with extracted ion chromatograms highlighting the ten most abundant deconvoluted intact IgG mass features. Corresponding deconvoluted mass spectra are shown in matching colors according to their retention-time windows. (B) Cosine similarity analysis of native intact IgG mass profiles between donors M54, F42, and F55, showing donor-specific intact IgG repertoire profiles under the applied matching criteria (results summarized in Table S1)

Across all three donors, native nCEC-MS separated the intact IgG repertoires over a broad chromatographic window. The observed elution behavior is consistent with charge-based separation of closely related IgG proteoforms, reflecting subtle differences in sequence, subclass, and other molecular features. Within the ten most abundant mass features of each donor, deconvoluted masses ranged from 143 to 149 kDa (Supplementary Figure 5), consistent with intact, N-glycan-truncated IgG species. Discrete proteoform signals were detected across most of the chromatographic separation, with regions exhibiting spectral convolution, resulting in no distinct spectra. In these regions, overlapping charge envelopes from highly complex mixtures of IgG species prevented reliable deconvolution, highlighting the analytical challenge posed by the dynamic range and molecular complexity of the circulating IgG repertoire.

The abundance of intact IgG mass features was unevenly distributed. Across the three donors, only 12-19% of the detected masses accounted for 50% of the cumulative relative abundance. We next applied cosine similarity to compare intact IgG mass profiles between donors (Figure 2B, values reported in Supplementary Table 2). Inter-donor comparisons, including M54-F42, F42-F55, and M54-F55, showed no detectable overlap among the identified intact mass features within the applied matching criteria (Supplementary Figure 5). This donor specificity agrees with previous studies of IgG1 repertoires at the domain level^5^.

### Middle-up profiling captures donor-specific Fab and Fc/2 diversity

To obtain orthogonal structural information complementary to the intact IgG mass profiles, the same donor samples were analyzed using middle-up RPLC-MS workflows targeting the Fc/2 constant region and the IgG1 Fab region.

Constant-region profiling was performed using IdeS digestion to generate single-chain Fc/2 subunits of approximately 25 kDa. The nRPLC-MS method was optimized to resolve and quantify Fc/2 subunits, including allotypes with nearly isobaric masses, providing information on subclasses, allotypes, and N-glycoforms (Figure 3B). Across all donors, IgG1 was the predominant subclass, accounting for 50-60% of the total Fc/2 signal. IgG2 was also abundant, contributing to 33–35% of the signal, while IgG3 (up to 7.5%) and IgG4 (5-12%) were less abundant. This diversity was expanded by the addition of N-glycans, resulting in 8-10 distinct glycoforms per allotype. Overall, monitoring Fc/2 masses revealed a highly diverse landscape of constant-region proteoforms, with more than 50 Fc/2 proteoforms per individual.

**Fig. 3:**
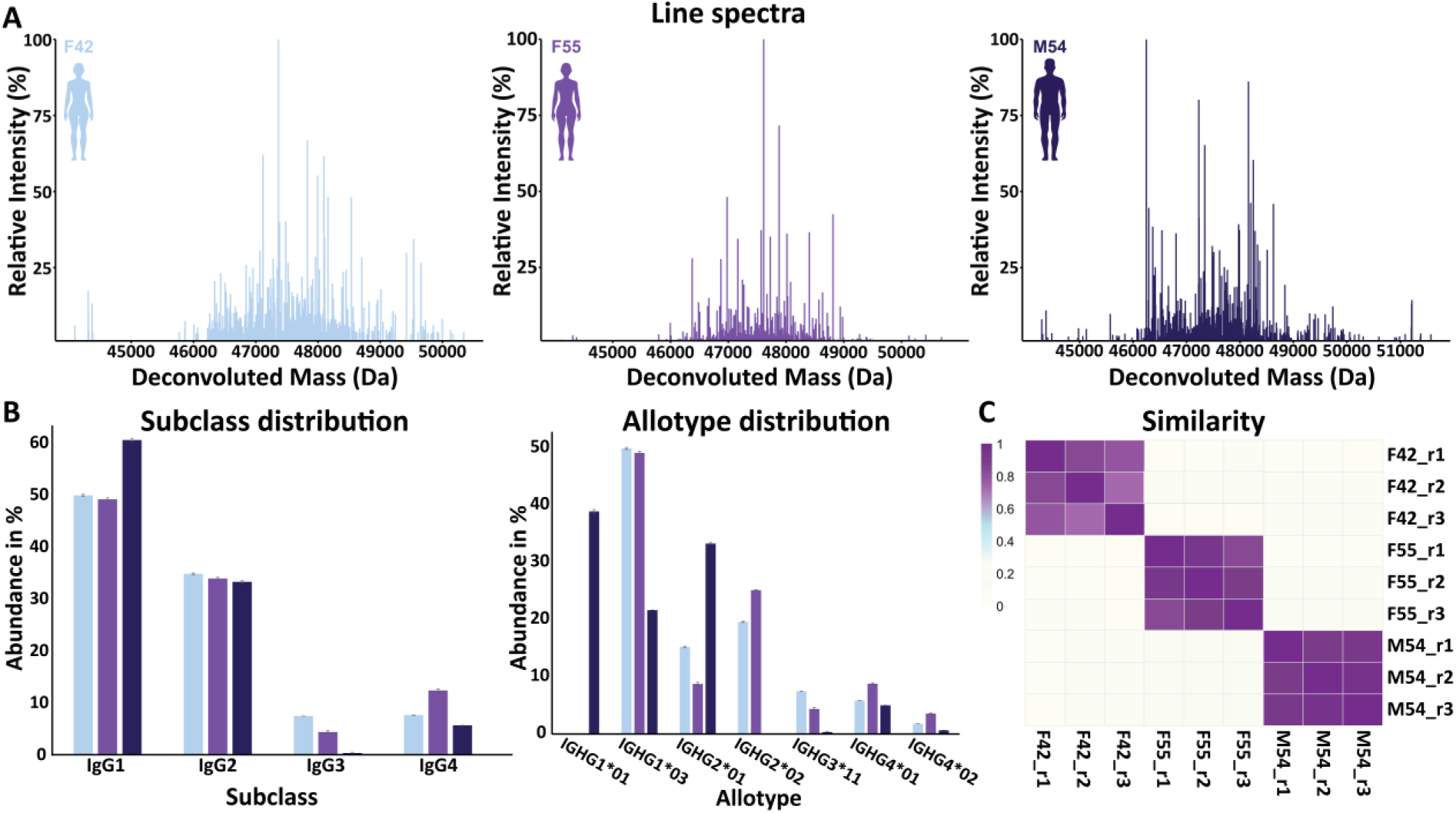
Middle-up profiling captures donor-specific Fab and Fc/2 diversity. (A) IgG1 Fab mass profiles of donors M54, F42, and F55 obtained after BdpK digestion and nRPLC-MS analysis. (B) Fc/2-based subclass and allotype distributions obtained after IdeS digestion, showing donor-specific constant-region profiles and the relative contribution of IgG subclasses. (C) Cosine similarity analysis of IgG1 Fab profiles across donors and replicates. The 200 most abundant Fab masses from each donor and replicate were compared using a 2 Da matching window, with color intensity indicating the degree of profile similarity (data summarized in Table S2).

In parallel, IgG1 Fab region profiling was performed after BdpK digestion. Given the selectivity of BdpK for IgG1, subsequent middle-up analysis was restricted to the IgG1 repertoire. BdpK digests IgG1 above the hinge region, generating intact Fab subunits of approximately 50 kDa. Using 0.2 min sliding chromatographic windows combined with maximum entropy deconvolution, we mapped the complex IgG1 Fab mass landscape.^22^ Across donors, approximately 500–800 distinct Fab masses were identified per individual in the *m/z* 1200–2400 range, with charge states between 21 and 33 (Figure 3A). As anticipated, middle-up analysis provided a considerably higher number of detectable mass features than intact IgG profiling. This higher number can be attributed to lower structural heterogeneity and lower mass of the Fab subunits, resulting in improved ionization efficiency and MS sensitivity. The profiles obtained aligned with the intact IgG results, revealing a heterogeneous variable-domain repertoire dominated by a smaller subset of high-abundance Fab mass features. Similarly to the intact IgGs, the obtained Fab masses were compared within a 2 Da window using cosine similarity. Replicate measurements showed high intra-donor similarity and low inter-donor similarity, supporting the reproducibility of the Fab profiling workflow and the donor-specific nature of the Fab repertoire.

The Fab abundance distributions were strongly skewed across all donors, with a minor fraction of abundant masses accounting for a large proportion of the cumulative abundance. Approximately 20% of the identified Fab masses accounted for 50% of the cumulative relative abundance across the three donors. This effect was particularly pronounced for donor F55, where only 9.5% of the identified masses accounted for 50% of the cumulative abundance. Conversely, looking at the low-abundant unique Fab masses, between 60% and 70% of the identified Fab masses accounted for 90% of the cumulative abundance, highlighting the extensive diversity of the Fab repertoire and its polyclonality. These observations are consistent with previous work by Bondt et al,^5^ who showed that the IgG clonal repertoire is highly heterogeneous but dominated by a subset of abundant clones.

### Integrated intact and subunit profiling reconstructs IgG molecular topology

To assess whether the native intact mass profiles were consistent with biologically plausible IgG structures, we integrated the intact and middle-up measurements. During the Fab profiling approach, only IgG1 were characterized, therefore, we focused on the reconstruction of these species. Additionally, IgG represented the largest fraction of the circulating IgG pool in all three donors, comprising 50-60% of the Fc/2 signal. Theoretical intact IgG1 masses were calculated by summing independently measured Fab masses with Fc/2 backbone masses carrying the remaining core GalNAc and fucose modifications. For donors F42 and F55, a single IGHG*03 allotype contributed to the IgG1 response, whereas donor M54 was heterozygous and exhibited two distinct IGHG1*01 and IGHG1*03 allotypes, both of which were used for reconstruction. As shown by the mirrored line spectra in Figure 4, the reconstructed IgG1 mass profiles aligned with the experimental native intact mass distributions measured by nCEC-MS. Due to the higher number of masses detected by the middle-up Fab approach, the reconstructed profiles resulted in increased density of the spectra profile. Still, the dominant signals showed a good alignment, providing cosine similarity scores between reconstructed and measured intact profiles of 0.30 for M54, 0.37 for F42, and 0.26 for F55 within-donors (Figure 4). In contrast, inter-donor similarities between reconstructed and measured intact profiles were below 0.15 for all comparisons. The detailed values are reported in Supplementary Table 1. It is important to note that the experimentally measured intact profiles represent the global antibody repertoire across all circulating IgG subclasses, whereas the reconstruction was restricted to IgG1 (which accounts for approximately 50% of the IgG repertoire) and was based on independently measured Fab and Fc/2 subunits. Despite this difference in molecular scope, the highest-intensity intact-mass features aligned with the predicted IgG1 clonal mass clusters. This supports the use of integrated native intact and middle-up profiling to partially reconstruct donor-specific IgG molecular topology by linking intact mass information with Fab diversity, subclass, allotype, and glycosylation features.

**Fig. 4:**
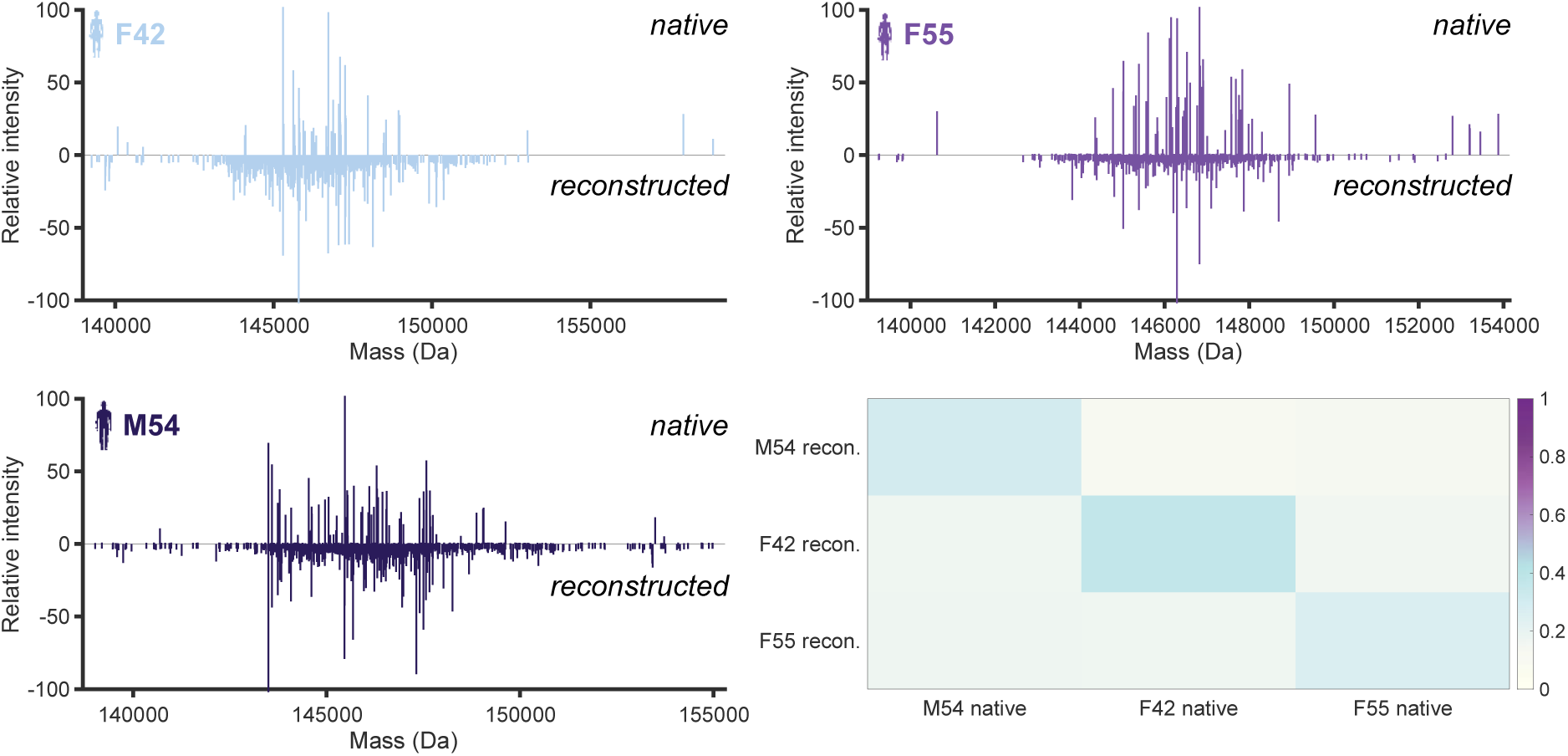
Integrated intact and subunit profiling reconstructs donor-specific IgG molecular topology. Mirrored line spectra comparing experimentally measured native intact IgG mass profiles consisting of IgG1-4 with reconstructed IgG1 mass profiles calculated from independently measured Fab and Fc/2 subunit masses. Reconstructed masses were calculated using IgG1 domain-level information, including the remaining core GalNAc and fucose modifications on Fc/2. Cosine similarity analysis comparing reconstructed (abbreviated as recon.) and experimentally measured intact profiles within and between donors, showing stronger alignment within matched donors than across unmatched donor comparisons (data summarized in Table S3).

## Discussion

Serum IgGs are among the most molecularly diverse proteins in circulation. This diversity arises from the combinatorial interplay of variable-region sequence variation, heavy- and light-chain pairing, subclass and allotype background, and Fc and Fab glycosylation. While state-of-the-art analytical strategies offer exceptional resolution for many of these features, they inevitably do so at decoupled molecular levels.^5,10^ Bottom-up approaches offer high compositional depth,^23,24^ while middle-up and middle-down approaches preserve distinct structural domains, such as isolated Fab and Fc subunits.^5,10^

However, both strategies physically disconnect these critical features from the intact antibody assembly. By hydrolyzing the IgG, these methods destroy the link between an individual clone’s antigen-binding identity (Fab) and its immunological potency (Fc). Consequently, determining which targeting domains coexist with a subclass-specific effector on a single circulating antibody remains an outstanding challenge.

This study presents a multi-level analytical framework for serum IgG repertoire analysis, bridging the gap by aligning intact IgG analysis with nCEX-MS and Fab and Fc profiling with middle-up nRPLC-MS. The main challenge in analyzing intact polyclonal IgG is the extreme molecular heterogeneity of the circulating repertoire. Without advanced online separation techniques, direct infusion native MS of purified serum IgG results in severe spectral congestion, in which hundreds of overlapping charge-state envelopes merge into an indistinct convoluted mass spectrum. Liquid-phase separation prior to MS is therefore essential to reduce sample complexity. While denaturing RPLC-MS is highly suitable for subunit separations (Fc and Fab), we found it to provide insufficient separation for intact endogenous polyclonal IgGs. Moreover, its acidic, organic mobile phases unfold antibody structure and distribute the ion current across a wide range (more than 10 charges) of highly charged states.

Our native nCEC-MS method addresses these limitations by combining charge-based chromatographic selectivity with near-native electrospray conditions, resulting in compact charge-state distributions (3 to 4 main charges) and improved spectral interpretability. Miniaturization to nanoflow conditions enabled intact IgG mass profiling from approximately 2 μg of purified IgG per analysis. Applied to three individual serum donors, this strategy detected 56-74 abundant intact IgG mass features per donor, with strongly donor-specific profiles. This extends previous observations of personalized antibody repertoires at the domain level^5^ by demonstrating that donor specificity is also preserved at the intact IgG level.

Our intact IgG profiles revealed that the circulating antibody landscape is not evenly distributed across the detected mass features, with a minor fraction of abundant intact masses accounting for the vast majority of the cumulative native signal. These findings are consistent with previous reports showing that the IgG repertoire is highly heterogeneous yet dominated by a subset of abundant clonal families. Currently, state-of-the-art Fab profiling relies on IgdE or BdpK enzymatic digestion, which are restricted to the IgG1 subclass.^25^ Our Fc analysis, however, confirms that IgG2, IgG3, and IgG4 collectively constitute 40-50% of the circulating pool across our donors. Consequently, middle-up workflows remain blind to nearly half of the functional antibody repertoire. In contrast, our nCEC-MS method does not suffer from this bias.

To confirm that our native intact signals represent IgG rather than artifacts from deconvolution or the gas phase, we used subunit-level measurements to rebuild the intact IgG clone masses in the donor samples. By adding together, the measured IgG1 Fab masses and their corresponding core-GalNAc \Fc backbones, we generated theoretical intact masses for each donor. Comparing these reconstructed masses with our experimental native nCEC-MS profiles within a 10 Da mass window revealed higher intra-donor cosine similarity scores: 0.297 (donor M54), 0.369 (donor F42), and 0.263 (donor F55). These values underscore a remarkable degree of similarity, given the differences in coverage (matched a single-subclass fragment platform in Fab profiling against a global IgG profile in intact screening), number of features (50-70 vs 600-800), and intensities. Their structural validity is supported by our inter-donor control results, in which the reconstructed masses of one individual were compared against the native intact spectra of an unrelated donor, yielding similarity scores of 0.106 to 0.173. This distinction proves that our integrated workflow successfully preserves structural features, demonstrating that the molecular topology information lost during offline domain-level hydrolysis can be recovered and verified.

Although these results highlight the value of moving towards intact IgG analysis and domain-level data integration, some limitations remain. First, the intact native measurements resolve the most abundant IgG mass features but do not yet capture the full depth of the circulating repertoire. Specific chromatographic regions still contain overlapping charge-state envelopes that could not be reliably deconvoluted, reflecting the large dynamic range and molecular complexity of serum IgG. Future implementations may benefit from combining high-resolution native nCEC separations with other liquid-phase (multidimensional HPLC) or gas-phase separations (ion mobility) or gas-phase charge-manipulation strategies (such as proton-transfer charge reduction) to further reduce spectral convolution and allow detection of lower-abundance IgG proteoforms.^26,27^ Second, the reconstruction presented here was restricted to IgG1, which was the dominant subclass in all three donors, but not the only contributor to the intact mass profiles. Extending this strategy to include IgG2, IgG3, and IgG4 Fab information would improve coverage of the total IgG repertoire. Third, EndoS2 digestion was used to reduce Fc glycan microheterogeneity and simplify intact mass interpretation. While this improves detection of the underlying antibody backbone, it also means that intact glycoform-level topology is not fully retained in the native measurement.

In conclusion, rather than seeking to replace established subunit-level workflows, the native-state platform presented here uses them as structural building blocks. By reconnecting Fab diversity with Fc subclass and allotype information using native intact mass profiling, the method provides a more integrated view of antibody heterogeneity and preserves molecular relationships that are otherwise lost.

Measuring this connectivity (molecular topology) is a prerequisite for transitioning from static lists of isolated sequence variants toward a complete, topological understanding of circulating IgG proteoforms and their precise functional roles in structural immunology, autoimmune diagnostics, and precision biotherapeutic design.

## Methods

### Serum samples and IgG concentration estimation

Serum samples from three individual healthy donors, M54, F42, and F55, were purchased from 3H Biomedical (Uppsala, Sweden). IgG concentrations were estimated using a NanoDrop spectrophotometer (Thermo Fisher Scientific) for both glycosylated and deglycosylated IgG samples. The estimated concentrations of glycosylated IgG were 0.697, 1.194, and 1.577 mg/mL for M54, F42, and F55, respectively. The corresponding concentrations of deglycosylated IgG were 0.671, 1.197, and 1.373 mg/mL, respectively.

### Sample preparation for intact IgG and Fc/2

IgGs were captured and eluted without on-bead hinge-region digestion as previously described by Blöchl et al.^10^ Fc N-glycans were hydrolyzed using EndoS2 endoglycosidase (Genovis, Lund, Sweden) at a ratio of 2 U/μg IgG. Digestion was performed for 30 min at 37 °C in 10 mM Tris, 150 mM NaCl, pH 7.4. EndoS2 cleaves native IgG Fc N-glycans after the core GalNAc, with or without core fucose. After EndoS2 digestion, samples were split into two fractions. The first fraction was analyzed directly by native MS for intact IgG profiling. The second fraction was subjected to below-hinge digestion using IdeS protease (Genovis, Lund, Sweden) at a ratio of 2 U/μg IgG in 20 μL of 150 mM ammonium bicarbonate, pH 7.6. Samples were incubated for 2.5 h at 37 °C to generate Fc/2 subunits^10^, and used to evaluate the efficiency of IgG deglycosylation. Glycosylated Fc/2 subunits were prepared from 1 μL serum or plasma of the same donors and characterized using a protocol developed by Blöchl et al^10^ .

The Fc/2 standard employed for system performance check was generated by digesting a commercial trastuzumab preparation (Herceptin; Roche Diagnostics GmbH, Penzberg, Germany) with IdeS at 1 U/μg IgG in 150 mM ammonium acetate (Sigma-Aldrich). The resulting Fc/2 standard was diluted in 150 mM ammonium acetate to a final concentration of 100 ng/μL.

### Sample preparation for IgG1-Fab analysis

For IgG1-Fab profiling, 10 μL of human serum or plasma from each donor was diluted to 100 μL in phosphate-buffered saline (PBS; 0.035 mM phosphate, 150 mM NaCl, pH 7.6) and spiked with 300 ng of trastuzumab standard (Herceptin; Roche Diagnostics GmbH). IgGs were purified using 10 μL of Fc-specific agarose beads (CaptureSelect™ FcXL Affinity Matrix; Thermo Fisher Scientific, Waltham, MA), self-packed into 96-well filter plates with 10 μm pores (Orochem Technologies, Naperville, IL). Samples were incubated for 2 h at room temperature with shaking at 900 rpm. The beads were washed twice with 200 μL PBS, twice with 50 mM Tris-HCl, 150 mM NaCl, pH 7.6, and once with 50 mM Tris-HCl, 150 mM NaCl, 10 mM CaCl₂, pH 7.6. After each washing step, the supernatant was removed by centrifugation at 500 × g for 2 min. IgG1 was then digested above the hinge region in a two-step process using BdpK protease (Genovis, Lund, Sweden) at a ratio of 2 U/μg IgG1 in a final volume of 30 μL. Each digestion step was performed for 1 h in a moisture box. After incubation, Fab subunits were collected by centrifugation at 500 × g for 2 min.

The Fab standard used for system performance check was generated by digesting commercial trastuzumab with BdpK at 1 U/μg IgG in 150 mM ammonium acetate (Sigma-Aldrich). The resulting Fab standard was diluted in 150 mM ammonium acetate to a final concentration of 500 ng/μL.

### Native nCEC-MS profiling of intact IgG

Intact IgG profiling was performed using nanoflow cation-exchange chromatography coupled to MS. The nano-cation exchange columns were prepared and packed in-house as previously described^19^. Samples were directly injected and separated on an in-house-packed nCEC column (100 μm i.d. × 150 mm, 5 μm, BioPro IEX SF resin, YMC) using a Vanquish Neo system (Thermo Fisher Scientific) coupled to an Orbitrap Ascend Tribrid mass spectrometer equipped with the BioPharma option (Thermo Fisher Scientific). The HPLC and MS systems were connected through a Nanospray Flex ion source (Thermo Fisher Scientific), a SimpleLink UNO connector (1/32; FossilionTech), and a hydrophobic-coated nano-emitter (20 μm i.d. × 75 mm; FossilionTech).

Separation was performed using mobile phase A, consisting of 50 mM ammonium acetate in H_2_O, pH 5.0, and mobile phase B, consisting of 250 mM ammonium acetate in H_2_O, pH 8.5. A 55 min gradient was applied at a flow rate of 500 nL/min with an injection volume of 1 μL directly on-column. The gradient started with a 2 min hold at 100% mobile phase A, followed by a linear increase to 100% mobile phase B over 30 min. The column was then held at 100% mobile phase B for 8 min, returned to 100% mobile phase A in 1 min, and re-equilibrated at 100% mobile phase A for 14 min.

MS data were acquired under native conditions with high-mass range mode activated. Data were recorded over an *m/z* range of 3000–10000 at a resolution of 17,500 at *m/z* 200. The spray voltage was set to 1.8 kV, the capillary temperature to 275 °C, the AGC target to 3.0 × 10^6^, the maximum injection time to 200 ms, and the number of microscans to 10. In-source CID was set to 85 eV.

### Intact IgG data analysis

Raw MS files were imported into FreeStyle v1.8 SP2 QF1 (Thermo Fisher Scientific). Mass spectra were manually extracted over the approximately 20 min IgG elution window by averaging spectra within 0.2 min windows, yielding about 100 averaged spectra per donor. The averaged spectra were imported into UniDec v7.0.1^28^ for deconvolution of intact IgG masses. Deconvolution was performed using an *m/z* range of 5500–7000 without background subtraction or signal normalization. A target mass range of 140–155 kDa was selected within a charge-state window of 20-28, with sampling every 0.1 *m/z*. Deconvoluted masses were picked every 10 Da using a threshold of 0.1 relative intensity. After deconvolution, spectra were manually classified, assessed, and filtered based on the mass-to-charge-state distribution and the DScore.^29^

The deconvoluted data were imported into MATLAB R2025b, where an in-house script was used for further analysis and plotting. Mass lists were normalized to the most intense mass feature for each donor, then filtered by manual classification. To assess similarity across donors, mass lists were binned using a bin width of 2 Da (Figure 2). For comparisons between measured and reconstructed intact profiles, a bin width of 10 Da was used (Figure 4). Cosine similarity scores were tshen calculated from the binned mass-intensity vectors. Extracted ion chromatograms were generated in FreeStyle by selecting three charge states per deconvoluted spectrum, applying 7-point Gaussian smoothing, and using a mass tolerance of 50 ppm.

### HPLC-MS profiling of Fc/2 and IgG1-Fab subunits

Fc/2 and IgG1-Fab were analyzed on an Ulitmate 3000 system (Thermo Fisher Scientific), equipped with a 5 uL loop. Deglycosylated Fc/2 were injected into a Vanquish Neo system (Thermo Fisher Scientific) equipped with a 20 μL sample loop to evaluate the deglycosylation process. For Fc/2 profiling, 2 μL of deglycosylated sample or 0.2 μL of glycosylated Fc/2 were injected, and samples were analyzed in duplicate. For IgG1-Fab profiling, 5 μL of sample and 1 μL of Fab standard were injected, and samples were analyzed in triplicate. Both workflows used a heated trap-and-elute configuration. Samples were loaded onto a C4 trap column (5.0 × 0.3 mm i.d., Acclaim™ PepMap™, 300 Å pore size; Thermo Fisher Scientific) at 60 °C and 15 μL/min under isocratic conditions using 0.1% trifluoroacetic acid (TFA; Merck, Darmstadt).

Chromatographic separation was performed on a diphenyl reversed-phase column (150.0 × 0.1 mm i.d., Halo® BioClass, 1000 Å pore size; Advanced Material Technology, Wilmington, DE) operated at 1 μL/min and maintained at 80 °C using an external column oven. Mobile phase A consisted of 0.1% TFA in H_2_O (ELGA Labwater, Ede, the Netherlands), and mobile phase B consisted of 0.1% TFA in acetonitrile (Actu-All Chemicals, Oss, the Netherlands). Fc/2 glycosylated subunits were separated similarly to Blöchl et. al. using a multi-step gradient from 20.0% to 27.8% B in 1.0 min, 27.8% to 32.0% B in 12 min, and 32% to 60.0% B in 2 min, followed by 90.0% B for 5 min and rapid column re-equilibration, while deglycosylated Fc/2 were analyzed similarly with a gradient between 30 and 33.5% B. IgG1-Fab subunits were separated using a 35-min gradient from 30.0% to 42.0% B, followed by an increase from 42.0% to 75.0% B in 5 min, 90.0% B for 4 min, and rapid re-equilibration at 20.0% B for 5 min.

The nanoHPLC system was coupled to a QTOF mass spectrometer (maXis Impact II, Bruker, Bremen, Germany) via a nano-ESI source (CaptiveSpray; Bruker), with acetonitrile-enriched gas applied at 0.6 bar using a nanoBooster system (Bruker). Measurements were performed in positive ion mode, using a source voltage of 1100–1350 V for Fc/2 profiling and 900 V for IgG1-Fab profiling. The source temperature was set to 220 °C, and the dry gas flow was set to 3.0 L/min. Collision cell RF was set to 2000 Vpp, the quadrupole ion energy and collision cell energy were set to 5.0 and 7.0 eV, respectively, and the transfer time was set to 150.0 μs. Funnel 1 RF and funnel 2 RF were set to 300.0 and 600.0 Vpp, respectively. In-source CID was set to 55 eV for Fc/2 profiling and 90 eV for IgG1-Fab profiling to provide sufficient declustering without fragmentation of the analytes. Data were acquired over an *m/z* range of 600-4000 with a rolling average of three scans at an acquisition rate of 1.0 Hz.

### Fc/2 and IgG1-Fab data analysis

Fc/2 data were analyzed using the workflow previously described by Blöchl et al.^10^. Briefly, initial allotype identification was based on protein average masses obtained by maximum entropy deconvolution of averaged raw mass spectra using Compass DataAnalysis software (Bruker). The resolution was set to 5000, and the data spacing was set to 1.0. Allotype identification and relative quantification were subsequently performed using Skyline v22.2.0.351 based on extracted ion current chromatograms. Before import into Skyline, raw spectra were cropped to 600-1320 s and *m/z* 1000-1600 to reduce data processing time and were converted to mzML files using OpenMS. Retention-time alignment was performed using LaCyTools. The efficiency of IgG deglycosylation was determined from the averaged deconvoluted mass spectra.

IgG1-Fab data were internally calibrated using the theoretical *m/z* values of the spiked trastuzumab Fab standard. Maximum entropy deconvolution was performed in Compass DataAnalysis software between 16 and 42 min using sliding 0.2 min windows. A threshold of 3000 absolute intensity and 0.1% relative intensity was applied within each window. The output mass range was set to 44000-52000 Da. Before deconvolution, each chromatographic window was baseline-subtracted and smoothed on default settings. Further data processing was performed in RStudio v4.3.3.1. Identified masses were filtered to remove mass shifts corresponding to glycation and oxidation, and masses derived from the spiked trastuzumab standard were removed. Final line spectra were generated using the masses that contribute to 90% of the total cumulative intensity. The identified Fab masses from all three donors were compared using cosine similarity with a 2 Da acceptance window.

## Data availability

The data needed to make all conclusions in this manuscript are present in the main text or can be found in the Supplementary Information. All raw spectra and HPLC-MS data from this study has been deposited to the Zenodo repository with identifier 10.5281/zenodo.21393355. All code used for figure generation and data analysis can be made available upon request.

## Acknowledgements

This research was funded by the Dutch Research Council (NWO) through the M2 HYPE-IMMUNe grant (OCENW.M.22.193). The work was conducted within the framework of the Chemometrics and Advanced Separations Team (CAST), and the authors gratefully acknowledge the contributions of its members. The authors thank Sabrina Reusch for her assistance with sample preparation for the Fc/2 measurements. The authors wish to thank Saar van der Laarse, Ales Holfeld, and Andrew Norris from Thermo Fisher Scientific for their support in enabling the measurements of the intact IgG data.

## Author contributions

TH, DM, CB, EDV, and AFGG conceptualized the research. Experimental work was performed by DM, ZZ, AAMvdZ, and CB. TH and DM performed the data analysis and interpretation and drafted the manuscript. AAMvdZ, ZZ, and CB critically reviewed the manuscript. EDV and AFGG acquired funding and contributed to writing the manuscript. All authors reviewed, edited, and approved the final version.

## Competing interests

The authors declare that they have no known competing financial interests or personal relationships that could have appeared to influence the work reported in this paper.

## Notes

### Competing Interest Statement

The authors have declared no competing interest.

