## Supplementary information for "Direct measurement and reconstruction of intact polyclonal IgG repertoires to preserve molecular and functional connectivity"

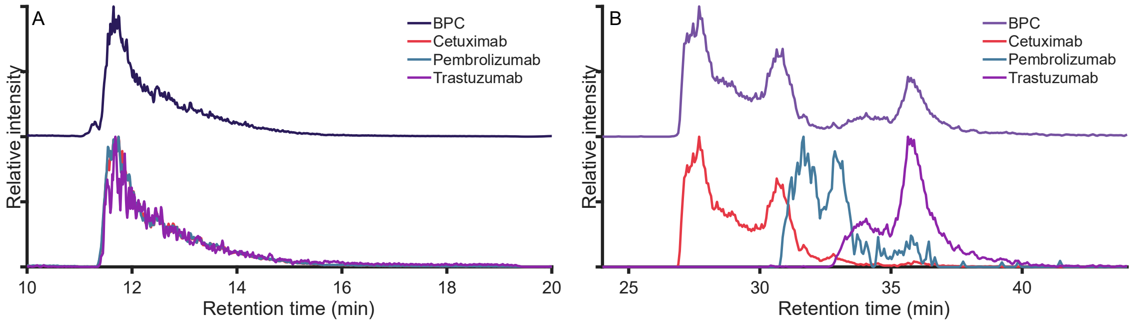


**Supplementary Fig. 1: Native nCEC-MS improves charge-based separation of standard monoclonal antibodies compared with nanoRPLC-MS.** (A) Base peak chromatogram and extracted ion chromatograms obtained by nanoRPLC-MS analysis of standard monoclonal antibodies. (B) Base peak chromatogram and extracted ion chromatograms obtained by native nCEC-MS analysis of the same monoclonal antibody mixture. All measurements were performed on a Q Exactive Plus mass spectrometer equipped with the BioPharma option.

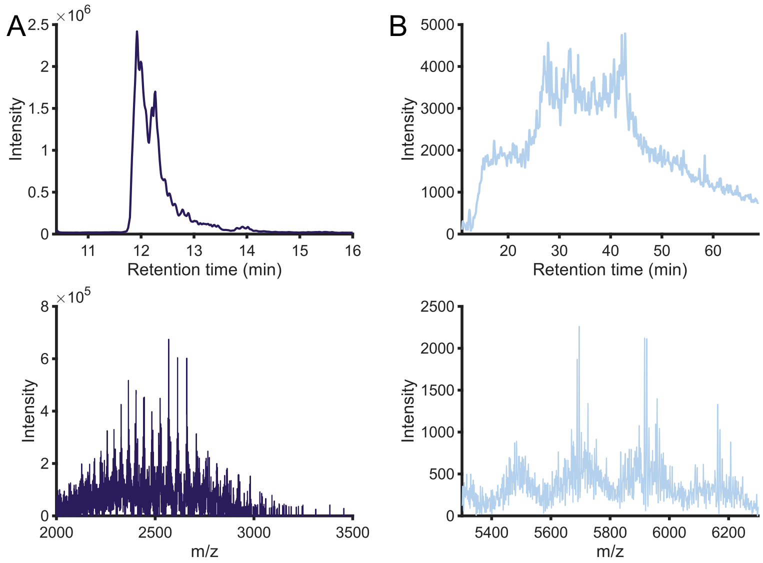


**Supplementary Fig. 2: Native nCEC-MS reduces spectral congestion during intact IgG profiling of donor M54.** (A) NanoRPLC-MS analysis of intact IgG from donor M54, with the corresponding mass spectrum obtained by averaging across the full IgG elution region. (B) Native nCEC-MS analysis of intact IgG from donor M54, with the corresponding mass spectrum extracted from a 0.2 min chromatographic window. All measurements were performed on a Q Exactive Plus mass spectrometer equipped with the BioPharma option.

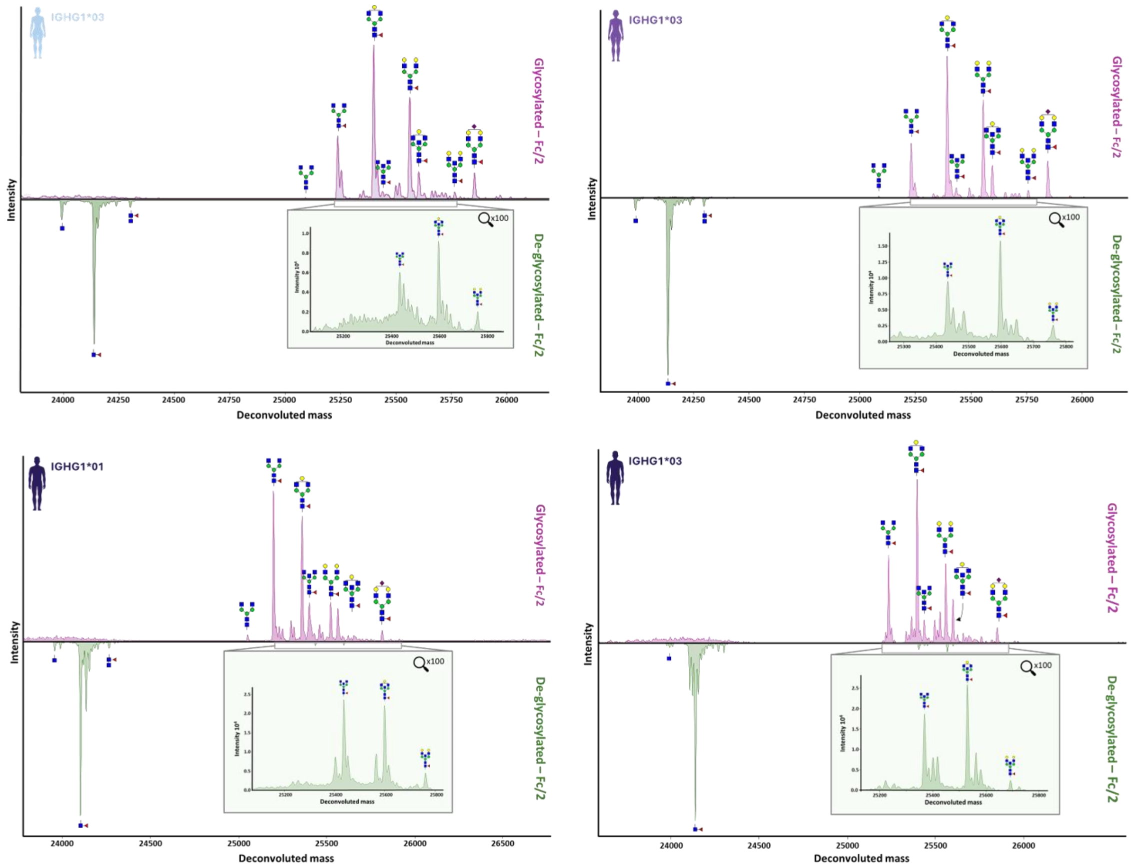


**Supplementary Fig. 3: EndoS2 digestion efficiently reduces Fc N-glycan heterogeneity in donor IgG samples.** Fc/2 subunits were analyzed before and after EndoS2-mediated deglycosylation to assess the removal of Fc N-glycan heterogeneity and confirm the generation of simplified Fc/2 mass profiles for downstream intact IgG interpretation.

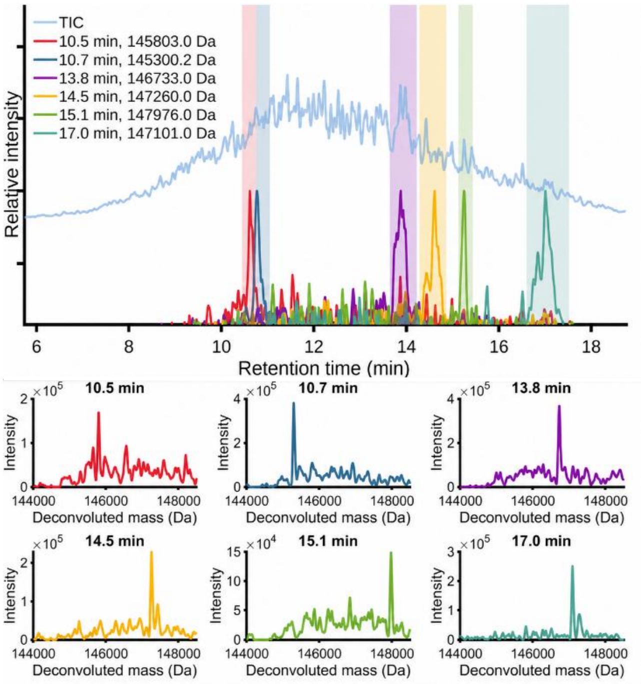


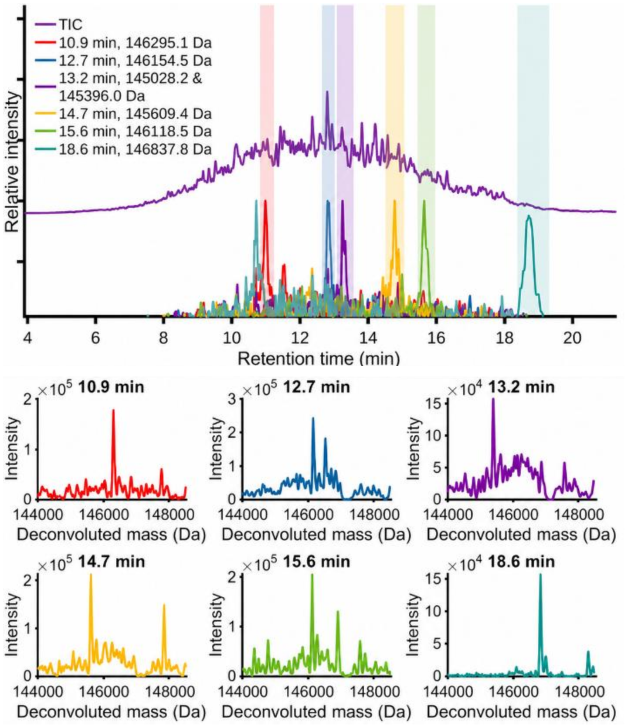


**Supplementary Fig. 4: Native nCEC-MS resolves donor-specific intact IgG mass features in donors F42 and F55.** Total ion chromatograms, extracted ion chromatograms, and corresponding deconvoluted mass spectra are shown for donors F42 and F55. Deconvoluted spectra are displayed in colors matching the selected chromatographic regions or extracted ion chromatograms.

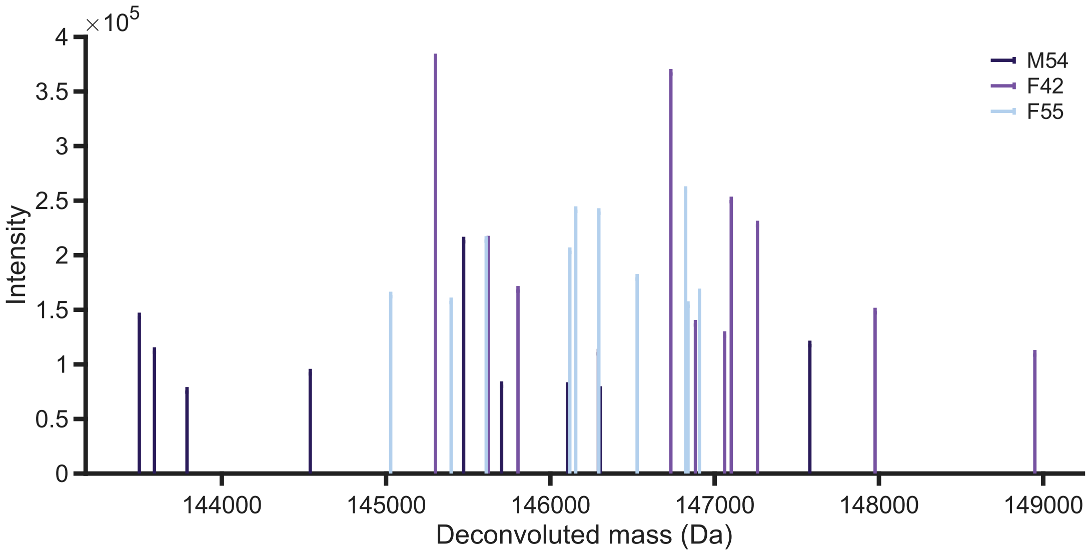


**Supplementary Fig. 5: The most abundant intact IgG mass features differ between donors.** The ten most abundant deconvoluted intact IgG masses are shown for each donor, highlighting the donor-specific nature of the abundant intact IgG mass profiles detected by native nCEC-MS.

**Supplementary Table 1:** Cosine similarity values for comparison inter-donor for native measurements and between reconstructed and native measurements. All masses were considered for the cosine similarity and a bin width of 10 Da.

| Native measurements | | | |
|---|---|---|---|
| donor | M54 | F42 | F55 |
| M54 | 1 | 0 | 0.07 |
| F42 | 0 | 1 | 0.03 |
| F55 | 0.07 | 0.03 | 1 |
| Native and reconstructed | | | |
| donor | M54 native | F42 native | F55 native |
| M54 recon. | 0.30 | 0.11 | 0.14 |
| F42 recon. | 0.17 | 0.37 | 0.16 |
| F55 recon. | 0.17 | 0.17 | 0.26 |

**Supplementary Table 2:** Cosine similarity values for comparison between the three single donors for IgG1 Fab middle-up measurements. The top 200 masses were used for the cosine similarity, with bins of 2 Da.

| Middle up IgG1 Fab measurements | | | | | | | | | |
|---|---|---|---|---|---|---|---|---|---|
| donor | M54_r1 | M54_r2 | M54_r3 | F42_r1 | F42_r2 | F42_r3 | F55_r1 | F55_r2 | F55_r3 |
| M54_r1 | 1 | 0.88 | 0.91 | 0.05 | 0.06 | 0.08 | 0.06 | 0.07 | 0.08 |
| M54_r2 | 0.88 | 1 | 0.95 | 0.05 | 0.07 | 0.06 | 0.06 | 0.07 | 0.06 |
| M54_r3 | 0.91 | 0.95 | 1 | 0.05 | 0.07 | 0.07 | 0.06 | 0.07 | 0.07 |

| | | | | | | | | | |
|---|---|---|---|---|---|---|---|---|---|
| F42_r1 | 0.05 | 0.05 | 0.05 | 1 | 0.80 | 0.75 | 0.04 | 0.05 | 0.06 |
| F42_r2 | 0.06 | 0.07 | 0.07 | 0.80 | 1 | 0.69 | 0.05 | 0.06 | 0.07 |
| F42_r3 | 0.08 | 0.06 | 0.07 | 0.75 | 0.69 | 1 | 0.03 | 0.04 | 0.05 |
| F55_r1 | 0.06 | 0.07 | 0.08 | 0.04 | 0.05 | 0.03 | 1 | 0.91 | 0.80 |
| F55_r2 | 0.07 | 0.07 | 0.07 | 0.05 | 0.06 | 0.04 | 0.91 | 1 | 0.87 |
| F55_r3 | 0.08 | 0.06 | 0.07 | 0.06 | 0.07 | 0.05 | 0.80 | 0.87 | 1 |

**Supplementary Table 3:** Number of identified IgG1 Fab masses in the three single donors and their contribution to the total repertoire.

| Identified masses | | | | | | |
|---|---|---|---|---|---|---|
| donor | replicate 1 | replicate 2 | replicate 3 | average | std | RSD% |
| M54 | 836 | 856 | 832 | **841** | 10.50 | 1.25 |
| F42 | 720 | 686 | 773 | **726** | 35.80 | 4.93 |
| F55 | 820 | 779 | 823 | **807** | 20.07 | 2.49 |
| **Cumulative 90 %** | | | | | | |
| donor | replicate 1 | replicate 2 | replicate 3 | average | std | RSD% |
| M54 | 544 | 560 | 540 | **548** | 8.64 | 1.58 |
| F42 | 532 | 506 | 542 | **527** | 15.17 | 2.88 |
| F55 | 497 | 477 | 497 | **490** | 9.43 | 1.92 |
| **The percentage of the total identified masses that represent 50%** | | | | | | |
| donor | replicate 1 | replicate 2 | replicate 3 | average | std | RSD% |
| M54 | 12.80 | 12.96 | 12.74 | **12.83** | 0.09 | 0.72 |
| F42 | 19.72 | 19.53 | 16.30 | **18.52** | 1.57 | 8.48 |
| F55 | 9.51 | 9.37 | 9.60 | **9.49** | 0.09 | 0.99 |
